# Neural mechanisms of lightness constancy

**DOI:** 10.64898/2026.11.19.767952

**Authors:** Fatemeh Didehvar, Robbe Vanstiphout, Patrick Cavanagh, Tom P. Franken

## Abstract

Object-specific variables are entangled in sensory input. For example, lightness, whether a surface is white or black, not only depends on the surface reflectance but also on the incident illumination. The brain must thus perform inference to achieve a reliable percept of lightness irrespective of illumination (lightness constancy). Whether this results from simple luminance contrast operations early in the visual system or computations at higher levels remains unresolved. We find that neural lightness signals only appear in the extragranular layers of cortical Area V4, and dominate for shadow regions, as opposed to the comparison of local and surrounding luminance (simultaneous contrast). This suggests that lightness is computed downstream from simultaneous contrast, ruling out simple low-level mechanisms, and instead supporting mid-level anchoring theory.

## Main Text

Perception is the cognitive act of constructing a representation of the external world, which enables us to efficiently interact with it (*1*). This is a challenging problem in all the senses, because real-world variables are entangled in the input that the senses receive (*1–8*). How the brain infers real-world variables remains poorly understood. This is true even for seemingly elementary cases, such as lightness, the perception of the reflectance of surfaces (whether a surface is black, gray or white) (*8–11*). Lightness is important for survival, as it aids in foraging and predation (e.g. indicated by countershading, one of the most common camouflage strategies, in which animals counterbalance natural patterns of illumination) (*12*, *13*).

The light that enters the eyes (luminance) is the product of reflectance and incident illumination. Because reflectance varies by a ratio up to ∼30, whereas incident illumination varies by a ratio up to ∼10^8^ (and is typically non-uniform), luminance poorly correlates with reflectance (*8*). A black surface in sunlight can thus easily reflect the same amount or more light than a white surface in shadow, yet lightness is approximately constant when illumination varies (lightness constancy) (*8–10*). Demonstrations such as Adelson’s checker shadow (**Fig. 1A**) (*8*, *10*, *14*, *15*) show that the visual system depends on contextual information to successfully infer lightness. Lightness is thus a particularly tractable case of perceptual inference, as it involves the inversion of a simple pixel-wise product.

**Fig. 1.**
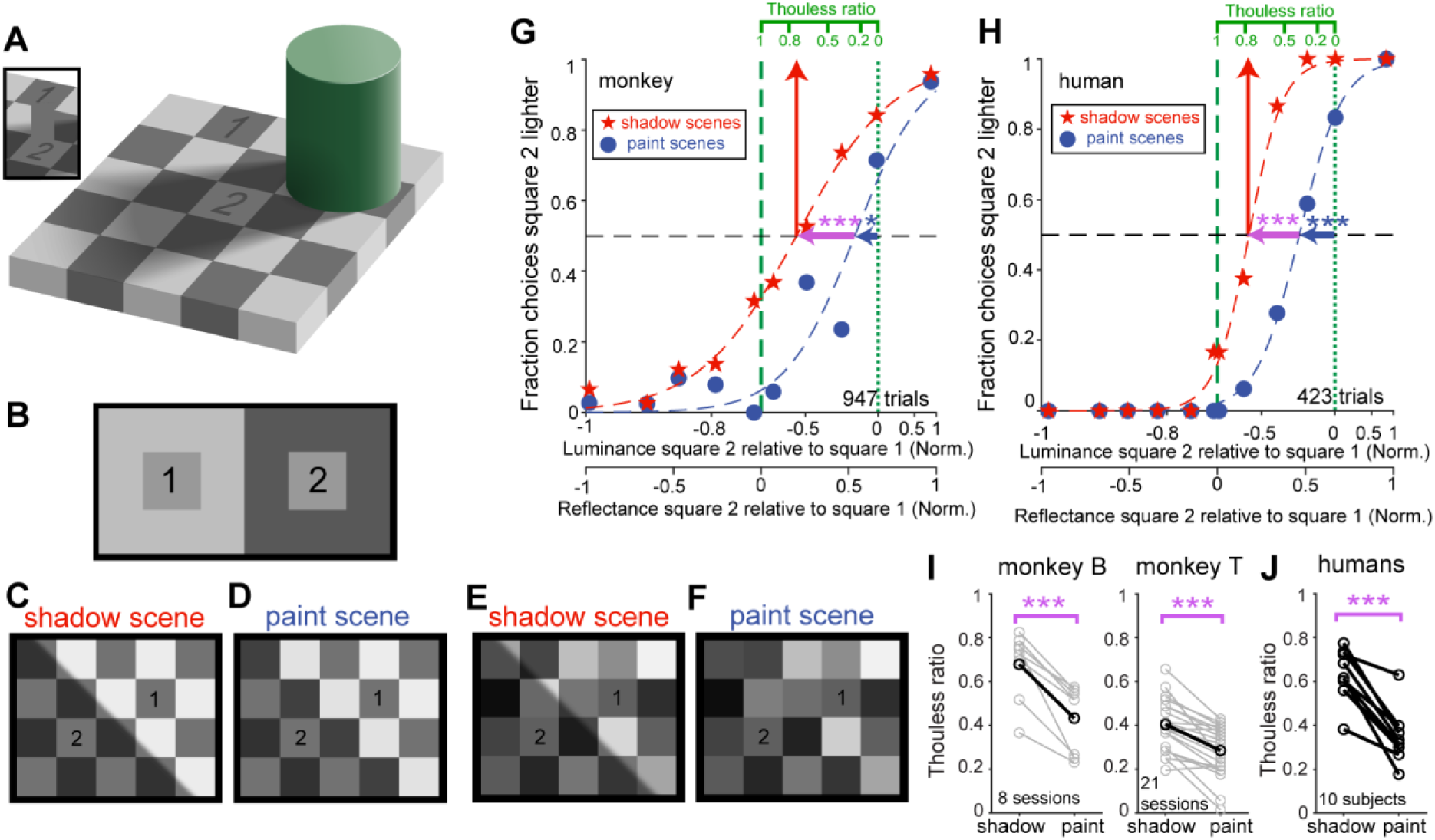
Macaques exhibit human-like lightness constancy. (**A**) Adelson’s checker shadow scene. Square 2 is perceived as lighter than square 1, even though their pixels are identical (see inset). (**B**) Simultaneous contrast. An identical gray square is perceived as lighter when the luminance of the surround is lower compared to when the luminance of the surround is higher. Squares 1 and 2 have the same pixel value. (**C**) Variant of A. (**D**) Paint: all pixels of each square in the scene in C are set to the average pixel value of that square, destroying the shadow border. (**E,F**) Example scenes used in the lightness task. Subjects reported which one of the two labeled squares appeared lighter. (**G**) Behavioral data from a macaque (example session). X-axis indicates the difference in luminance (reflectance) between square 2 and square 1, divided by their sum. Dashed red and blue lines: logistic function fits. (**H**) Similar to G, for data from an example human subject. (**I,J**) Thouless ratios for both monkeys (I; gray: individual sessions; black: average across sessions) and humans (J). ***: p<0.001; *: p<0.05.

Several theories of how lightness may be computed have been proposed. Low-level theories suggest that lightness results from simple spatial filtering operations, such as lateral inhibition, early in the visual system (*16–19*). Such theories can explain the perception of simultaneous contrast: two neurons whose classical receptive fields (cRFs) are positioned in the inner squares of **Fig. 1B** would receive a different amount of inhibition from neurons whose cRFs are positioned in the regions surrounding the inner squares, resulting in a different response. In contrast, mid- and high-level theories propose that lightness computations occur at higher stages of the visual system, which explicitly discount the illumination pattern (*3*, *10*, *20–24*).

To distinguish these theories, one needs to use scenes with spatially varying illumination, because these theories make different predictions for such scenes (*8–10*, *25*). In contrast, nearly all neuroscience studies have used variations of simultaneous contrast. Several of those studies identified neurons in primary visual cortex (V1) whose preference for luminance within their cRF is consistent with their firing to scenes in which only the luminance of the surround is manipulated (*26–30*). However, these neurons were small in number and their responses often highly heterogeneous, casting doubt on whether the net representation by a population of neurons was consistent with perception. Moreover, perceptual phenomena suggest that lightness is more than computing simultaneous contrast (*10*, *11*, *31*, *32*). For example, averaging the pixels of each square in the checker shadow scene (resulting in ‘paint scenes’) strongly reduces the lightness difference, even though simultaneous contrast remains the same (**Fig. 1C,D**) (*33*). One neurophysiology study that used spatially varying illumination did not identify a clear neural correlate for lightness (*34*) (see also Discussion).

Here we address this gap through behavioral experiments and Neuropixels recordings in macaque monkeys. Our behavioral experiments showed that macaques have human-like lightness constancy. We then used Neuropixels probes to record from large populations of neurons in macaque visual cortical Area V4 while presenting scenes with spatially varying illumination. Our recording approach samples simultaneously recorded populations in all laminar compartments. We find that a significant fraction of columnar populations of neurons exhibit net lightness signals. Interestingly, these signals do not occur in the granular (input) layer, and dominate in shadows, as opposed to the neural representation of simultaneous contrast. The neural lightness bias in V4 is sufficient to explain the behavioral lightness bias that we measured in these animals. Together, our data indicate that lightness is computed downstream from simultaneous contrast, ruling out simple low-level mechanisms, and instead supporting mid-level theories.

## Results

### Macaques exhibit human-like lightness constancy

We first wondered how the perceptual constancy for lightness of macaques compares to that of humans. Previous studies did not quantify lightness constancy in macaques. The lack of a clear neural correlate with lightness in a recent study led to the suggestion that macaques might not experience the checker shadow effect (*34*). To tackle this issue, we developed a lightness task for macaques. We presented scenes as in **Fig. 1E,F** while the macaque fixated the center of the display. The animals were trained to report with an eye movement whether square 1 or square 2 was higher in luminance (Materials and Methods). We used scenes with a pseudorandom distribution of reflectance values, to avoid that the animal would rely on the luminance contrast between neighboring squares to perform the task. We varied scene type (shadow scenes or paint scenes), and the luminance contrast between the squares of interest. We found that the psychometric function for shadow scenes was shifted leftward relative to that of the paint scene trials (magenta arrow in **Fig. 1G**, logistic regression model p-value for scene type: p = 2.2 × 10^-4^). The animal is thus more likely to report that a given square is the lighter one if it is located in a shadow as compared to an identical square in the corresponding region of a paint scene, consistent with the checker shadow effect. We quantified this shift using the Thouless ratio (*τ*), a standard constancy index (*8*, *35*, *36*). If vision would completely discount the illumination pattern (perfect lightness constancy), the shift would center the psychometric function on the vertical dashed line in **Fig. 1G** (*τ* = 1). If lightness constancy was fully absent (perfect luminance report), the psychometric function would be centered at the vertical dotted line (*τ* = 0). *τ* was significantly higher for shadow scenes than for paint scenes across sessions in both macaques (**Fig. 1I**; monkey B: 0.68 and 0.43, respectively, paired-sample t-test t_7_ = 6.0, p = 5.2×10^-4^; monkey T: respectively 0.40 and 0.29; t_20_ = 6.46, p = 2.7×10^-6^). These data demonstrate that macaques experience the checker shadow effect.

We observed that the psychometric function for paint scenes was also shifted left relative to a perfect report of luminance (bootstrap test p<0.05). Across sessions, *τ* for paint scenes was significantly larger than 0 for both monkeys (monkey B: one-sample t test t_7_ = 7.9, p = 9.3×10^-5^; monkey T: t_20_ = 11.1, p = 5.5×10^-10^). These data indicate that there is a perceptual simultaneous contrast effect in paint scenes, as the squares that surround square 2 are on average lower in luminance than those that surround square 1. Indeed, when we randomly repositioned all squares except for the pair that the animal is comparing, the psychometric function shifted to the right (fig. S1).

We then quantified lightness constancy in naïve human subjects using the same type of scenes. **Fig. 1H** shows data from an example human subject. As expected, and similarly to macaques, the psychometric function for shadow scenes was significantly shifted to the left compared to paint scenes (logistic regression model p-value for scene type: p = 4.1 × 10^-7^). In addition, the psychometric function for paint scenes was shifted left relative to a perfect report of luminance (bootstrap test p<0.001). These findings were confirmed in the population of human subjects (**Fig. 1J**): *τ* for shadow scenes was significantly higher than for paint scenes (paired-sample t-test t_9_ = 7.5, p = 3.6×10^-5^), and *τ* for paint scenes was significantly larger than 0 (one-sample t test t_9_ = 9.4, p = 5.9×10^-6^). These data are consistent with prior findings that lightness constancy in humans is incomplete (*τ* <1) (*8*). Together, our data show that macaques exhibit significant and human-like lightness constancy in checker shadow scenes, and that this constancy consists both of a component of discounting the illumination pattern, and of a component of simultaneous contrast. Next, we investigated the neural correlate of these components.

### Cortical Area V4 contains an inherent lightness bias

We used Neuropixels probes to record from columnar populations of neurons in cortical Area V4 in macaques. We used a recording chamber in which we replaced the native dura with a thin, transparent artificial dura, which enabled us to easily insert the thin Neuropixels probes into dorsal V4 while avoiding pial vessels (**Fig. 2A**) (*37–39*). We inserted the probes vertically such that the aggregate classical receptive fields of neurons recorded in the same session overlapped in visual space (**Fig. 2B**). We presented shadow scenes, similar to Adelson’s checker shadow, such that the receptive fields were centered on one square in the shadow region, and corresponding paint scenes (**Fig. 2C**). We also included shuffled scenes, which were derived from paint scenes by randomly reordering the other squares of the scene, to remove the resulting simultaneous contrast effect that exists in those scenes. We varied square luminance, and each scene was shown for 500-800 ms while the animal was fixating.

**Fig. 2.**
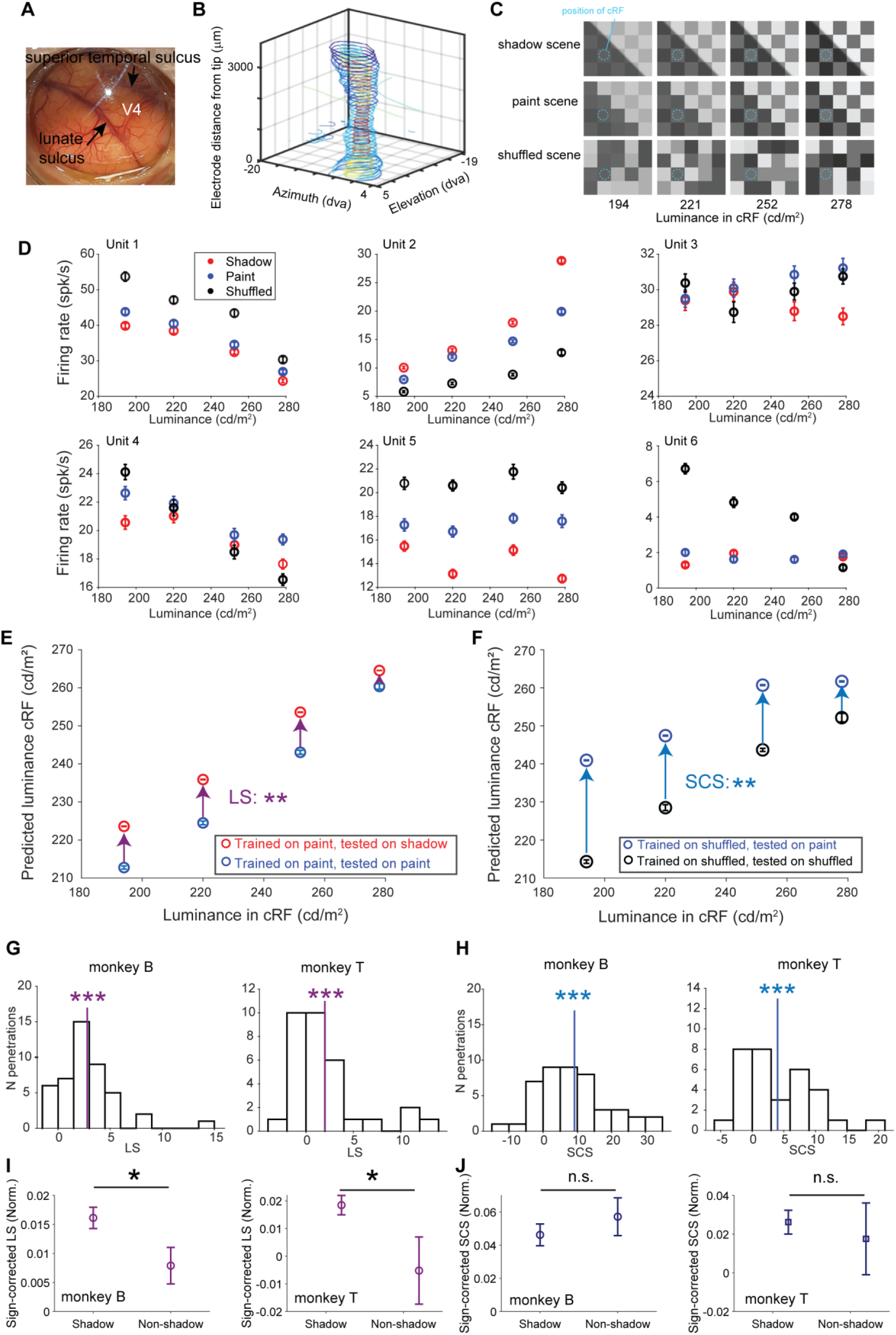
Lightness signals in cortical Area V4. (**A**) Artificial dura recording chamber positioned over cortical Area V4. (**B**) Aggregate classical receptive field (cRF) on each electrode, for an example recording session (see Methods). (**C**) Overview of stimuli. (**D**) Response of six units to the scenes shown in C. (**E**) Predicted luminance values from between-condition decoding analysis, for an example penetration. Lightness signal (LS) is defined as the difference in predicted luminance between shadow and paint scenes. (**F**) Similar to E but for the simultaneous contrast signal (SCS), defined as the difference in predicted luminance between paint and shuffled scenes. (**G**) Histogram of LS for both animals in all penetrations. Vertical line: average LS across penetrations. (**H**) Similar to G but for SCS. (**I,J**) Comparison of lightness signal (I) and simultaneous contrast signal (J) for conditions in which the cRFs were positioned either in a square in the dark (shadow) region or in a square in the highly illuminated (non-shadow) region of the scene (or the corresponding regions in paint or shuffled scenes). Signals are normalized to the luminance of the square with the cRFs, and sign-corrected. Signals for shadow and non-shadow were statistically compared with a paired t-test. Data in panels D, E, I and J plotted as mean ± s.e.m. ***: p<0.001; **: p<0.01; *: p<0.05. n.s.: not significant.

**Fig. 2D** and fig. S2 show data from six example single units. Unit 1 fires more to low luminances, irrespective of scene type (two-way ANOVA F(3,474) = 71.5, p = 4.4 × 10^-38^). In addition, the unit’s response also differed significantly between scene types (F(2,474) = 39.2, p=1.8×10^-16^). It fires less to paint scenes than to shuffled scenes with the same luminance. This preference is thus consistent with simultaneous contrast, given the unit’s preference for low luminance: it responds to a paint scene as if the luminance was higher than for a shuffled scene with the same luminance. Interestingly, the unit also fires differently to shadow and paint scenes: it responds to a shadow scene as if its luminance in the cRF was higher than that of a paint scene with identical luminance. This difference is thus consistent with the checker shadow effect. Unit 2 was also selective for luminance, but prefers high luminances (F(3,890) = 244.39, p = 1.7×10^-115^). Now the difference between scene types is reversed: the unit fires less to shuffled scenes than to paint scenes, and less to paint scenes than to shadow scenes (F(2,890) = 221.63, p = 9.4×10^-79^). This pattern is thus, again, consistent with perception. That said, for many units the patterns were harder to interpret. The response to luminance was sometimes non-monotonic (e.g. Unit 3), there was an interaction between the tuning for scene type and for luminance (e.g. Units 4,6), or the response differed for scene type but not for luminance (e.g. Unit 5). There is thus a strong heterogeneity in how V4 neurons respond to isoluminant patches, which is consistent with prior studies that used simultaneous contrast scenes (*26–30*).

Because of this heterogeneity it is not clear whether there is a net lightness signal. To tackle this question, we used a between-condition decoding approach to investigate whether the most accurate representation of luminance in the population of neurons is biased towards lightness (fig. S3). We trained random forest regressors to decode the luminance of the square with the cRFs, based on the spike counts from all simultaneously recorded units, from 90% of the trials with paint scenes. We then tested these decoders separately on responses from the remaining 10% of trials with paint scenes, and on responses from trials with shadow scenes. **Fig. 2E** shows the result for an example penetration. The predicted luminance is significantly higher when testing on shadow scene data than when testing on paint scene data (permutation test p≤0.01). The neural representation of luminance is thus biased towards higher values for shadow data than for paint data, consistent with our perception of such scenes (**Fig. 1C**). We define the average difference between the predicted luminance for shadow data and paint data as the neural lightness signal (LS; purple arrow in **Fig. 2E**). The average LS across penetrations was significantly positive, in both macaques (**Fig. 2G**; monkey B = 2.7, one-sample t-test t_44_ = 6.5, p=7.3×10^-8^; monkey T = 2.1, t_31_ = 3.8, p=6.8×10^-4^). We find similar results if the scene type used during testing was always different from the scene type used during training (fig. S4). The most accurate representation of luminance in columnar populations of V4 neurons is thus inherently biased towards lightness.

We can quantify the neural simultaneous contrast signal (SCS) in a similar way, by comparing responses to paint scenes and shuffled scenes. **Fig. 2F** shows the result of an example session, with a significantly positive SCS (permutation test p<0.01). The average SCS across sessions was also significantly larger than zero in both animals (**Fig. 2H**; monkey B=7.6, one-sample t-test t_44_ = 5.5, p = 1.9×10^-6^; monkey T=4.1, t_31_ = 4.1, p = 2.5×10^-4^). Simultaneously recorded populations of V4 neurons thus contain a net simultaneous contrast signal that is consistent with perception.

### Lightness signal dominates in shadow regions, as opposed to simultaneous contrast

The data that we presented thus far were derived from neurons with cRFs in the shadow region, but the checker shadow effect might also depend on discounting the brighter illumination in the lit region. To investigate that, we compared neural populations whose cRFs were either centered on a square in the shadow region or on a square in the illuminated region of the shadow scenes, or on corresponding squares in paint scenes and shuffled scenes. This analysis revealed that LS is strikingly asymmetrical: for both monkeys its magnitude is significantly larger in the shadow region than in the non-shadow region (**Fig. 2I**; paired t-test p = 0.0045 for data in both monkeys combined, 26 penetrations in monkey B and 14 penetrations in monkey T). This is in contrast to the SCS, which has a similar magnitude in both illumination regions (**Fig. 2J**). These data thus indicate that, whereas dark and bright regions contribute similarly to simultaneous contrast, the discounting of illumination dominates in shadow regions.

### No net lightness signal in granular layer, as opposed to simultaneous contrast

Low-level theories propose that lightness is computed early in the visual system, leading to the prediction that V4 inherits lightness signals from upstream areas such as V1 or V2. Feedforward inputs to V4 primarily target the granular (input) layer, as opposed to recurrent pathways (cortico-cortical feedback, intra-areal horizontal connections), which avoid the granular layer (*40–45*). The laminar organization of LS in V4 is thus informative regarding the pathways that are involved in the computation of lightness. If simple spatial filtering operations early in the visual system compute lightness, LS should already be strong in the granular layer of V4. To identify the position of laminar compartments in our data, we applied current source density (CSD) analysis as in prior studies, estimating the position of the granular layer as the current sink with the shortest latency (**Fig. 3A**) (*37–39*, *46–51*). We found a striking difference in LS between laminar compartments: LS was only significantly larger than zero in extragranular compartments, in both monkeys (**Fig. 3B**). The LS was weak in the input layer irrespective of the number of units included in the analysis (fig. S5). In contrast to LS, SCS was strong in the granular layer, for both monkeys (**Fig. 3C**). The latter is consistent with prior studies that found that neurons already in V1 signal simultaneous contrast when their receptive field is positioned within an isoluminant patch (*26–30*). Together, our data argue against the notion that LS in V4 is inherited from upstream cortical areas.

**Fig. 3.**
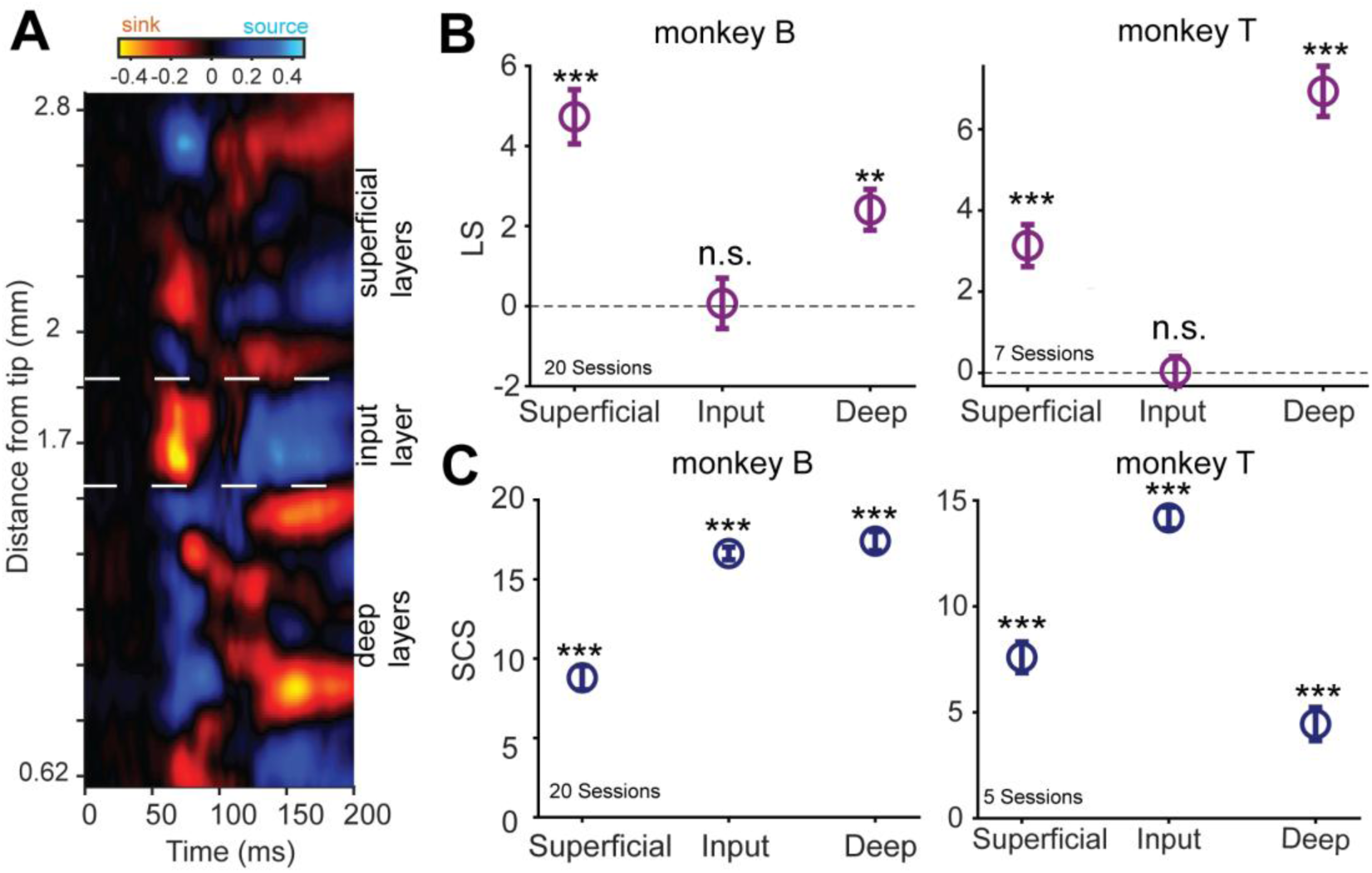
Lightness signal is weak in input layer, as opposed to simultaneous contrast signal. (**A**) CSD map for example Neuropixels recording. (**B**) Lightness signal (LS) in each laminar compartment. Mean ± s.e.m. Permutation test, monkey B: p_Superficial_ ≤ 0.001, p_Input_ = 0.46, p_Deep_ = 0.002; monkey T: p_Superficial_ ≤ 0.001, p_Input_ = 0.49, p_Deep_ ≤ 0.001. n.s.: not significant. (**C**) Similar to B but for simultaneous contrast signal (SCS). Permutation test, monkey B: p_Superficial_ ≤ 0.001, p_Input_ ≤ 0.001, p_Deep_ ≤ 0.001, monkey T: p_Superficial_ ≤ 0.001, p_Input_ ≤ 0.001, p_Deep_ ≤ 0.001.

### Lightness is computed later than simultaneous contrast

The laminar organization that we uncovered suggests that the computation of lightness depends on recurrent pathways in V4, as opposed to simultaneous contrast, which is computed as early as V1. If so, the difference between shadow and paint scenes is expected to be signaled later than the difference between paint and shuffled scenes. To test this prediction, we used demixed principal component analysis (dPCA) to measure the dynamics of the component of the population neural activity that differentiates either shadow scenes from paint scenes, or paint scenes from shuffled scenes (*52*). dPCA allows us to summarize the main features of neural activity even if firing patterns of individual neurons are heterogeneous. The first scene-type component is nearly independent of luminance, indicating that dPCA worked well (**Fig. 4A**). We find a latency of the context difference between shadow and paint scenes of 59.0 ms in both monkeys. In contrast, the latency of the difference between paint scenes and shuffled scenes was shorter, 41.1 ms in monkey B and 32.1 ms in monkey T (**Fig. 4B**). The difference between paint and shuffled scenes arises thus approximately 20 ms earlier than that between shadow scenes and paint scenes. To compare the latency of LS and SCS, we performed a sliding window between-condition decoding analysis (Methods). This revealed that the latency of SCS is shorter than that of LS (**Fig. 4C,D**).

**Fig. 4.**
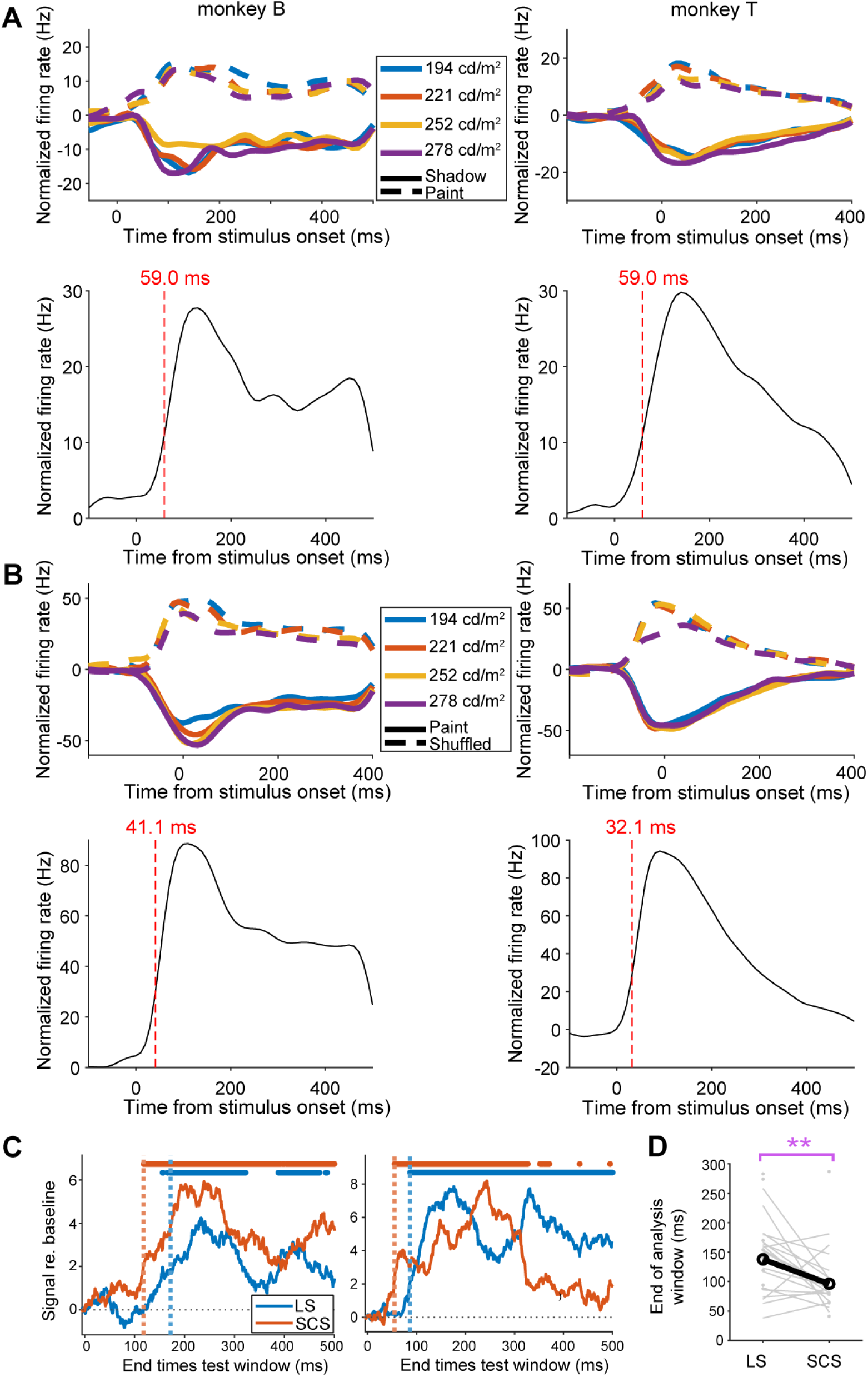
Dynamics of lightness signal. (**A**) Top: first component of dPCA that differentiates shadow scenes from paint scenes. Colors indicate different luminance values of the square with the cRFs. Bottom: difference curve between the data for the shadow scenes and the data for the paint scenes in the top panels. Vertical dashed line indicates the latency (threshold 33% of the peak re. baseline). Each column shows data from one monkey. (**B**) Similar to A but for the component that differentiates paint scenes from shuffled scenes. (**C**) Time course of LS and SCS, computed as between-condition decoding by random forest regressors in a sliding test window (100 ms wide). Symbols at the top show for which test windows the signal was significantly larger than 0. Latency was defined as the first of 15 consecutive windows that reached statistical significance. Each panel shows data from one penetration (left: monkey B; right: monkey T). (**D**) Population data for latency of LS and SCS. Gray lines: sessions for which latency was computable for both LS and SCS (paired). Gray dots: sessions for which latency was computable for only LS or SCS (unpaired). Black: average across paired data. **: sign rank test p = 0.009.

### Columnar V4 populations can account for perceptual lightness constancy

Next we asked to what extent the net neural lightness signal in individual columnar populations of V4 neurons is sufficient to explain the behavioral lightness constancy in these animals (**Fig. 1**). We derived neural Thouless ratios (nTR) from the penetrations in which we found significant LS and SCS, as we wondered to what extent the columnar populations with the strongest lightness signals compare to perception. We analyzed units in extragranular compartments, as we found that the lightness signal dominates in those compartments. We first trained random forest regressors to decode the luminance in square 2 relative to square 1 from responses to shuffled scenes. We then obtained the luminance predicted by these regressors for responses to shadow scenes. We obtained neurometric functions by computing the probability that the predicted luminance of square 2 relative to square 1 was positive. **Fig. 5A** shows the result of this analysis for one session. We can compute nTR for these data, similar to the behavioral data. We obtained nTR that are large enough to explain the behavioral TR (**Fig. 5B**). Together these data suggest that V4 contains columnar populations of neurons in which the representation of luminance is sufficiently shifted to account for perception.

**Fig. 5.**
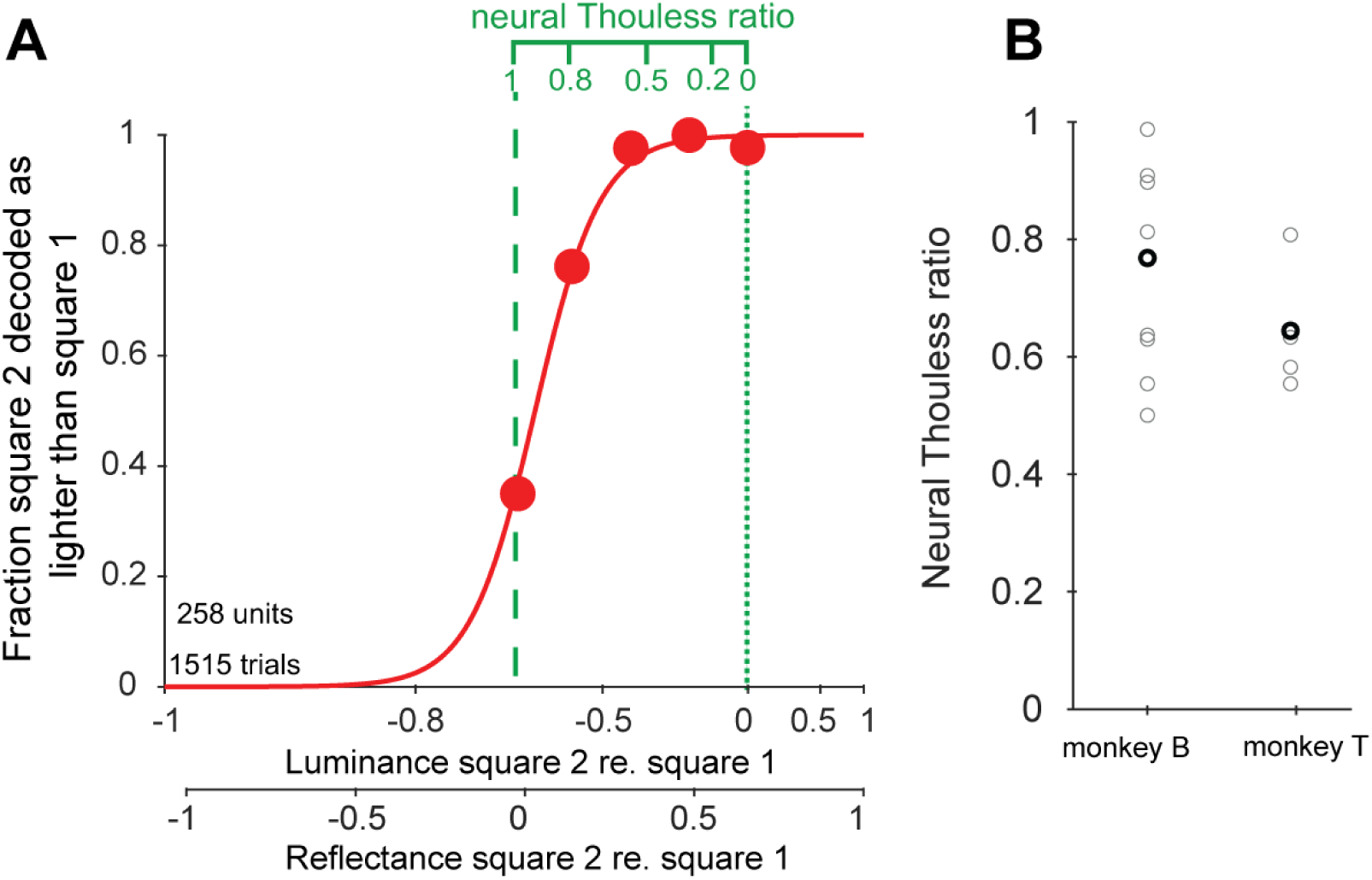
Columnar V4 populations can account for perceptual lightness constancy. (**A**) Fraction of trials for which the square in the shadow is predicted to have a higher luminance than the comparison square in the illumination region. X-axis indicates the luminance of the square in the shadow relative to that of the comparison square (as in Fig. 1G). Red line: logistic regression fit. (**B**) Neural Thouless ratios (derived from the analysis in A) for monkeys B and T (gray: individual sessions; black: average across sessions).

## Discussion

Our data advance insight into the neural mechanisms of how the brain computes lightness, a fundamental quality of vision. We establish macaques as a model to study this question, finding that these animals exhibit human-like lightness constancy. Our Neuropixels recordings in macaque visual cortex identify a net lightness bias in how columnar populations of V4 neurons represent the luminance of a surface. The contextual shift in luminance representation by these neural populations was sufficient to explain behavioral lightness constancy. Several findings suggest that the neural computation of lightness does not result from simple spatial filtering operations early in the visual system, as opposed to simultaneous contrast. First, the neural representation of lightness is delayed relative to that of simultaneous contrast. Second, the representation of lightness is weak in the input layer of V4, as opposed to the representation of simultaneous contrast. Third, the computation of lightness dominates in shadow regions, as opposed to the computation of simultaneous contrast. This indicates that lightness is computed downstream from simultaneous contrast.

Several studies identified neurons, as early as V1, that are sensitive to the ratio of the luminance in their cRF and the surrounding luminance (*26–30*). We find SCS in the granular layer of V4, consistent with a model in which this signal is inherited from upstream areas. That said, our finding that the context information necessary to compute SCS appears at ∼40 ms in V4, whereas many V1 units sensitive to luminance ratios have latencies >100 ms (*30*), suggests that these signals in V1 may also in part depend on feedback from V4.

The strong difference in LS between laminar compartments provides an explanation for why an earlier study did not identify a clear neural correlate for the checker shadow effect in either V1 or V4 (*34*). The authors discussed a few possibilities of why this may be the case. One possibility raised was that monkeys do not experience the paint-shadow effect in perception. We ruled this out in our behavioral experiment. Another possibility was that they did not record from the cortical layers in which the most relevant neurons are found. They used chronically implanted electrode arrays with a 1-mm electrode length, thus most of their neurons were likely positioned in the middle cortical layers. Using Neuropixels allowed us to record from a large number of neurons in each penetration, from all cortical layers. Our laminar analysis indicates that this is indeed a valid explanation: we do not find a clear lightness signal in the granular layer of V4, as opposed to superficial and deep cortical layers. Because feedforward inputs preferentially target the granular layer whereas recurrent and feedback pathways preferentially involve superficial and deep layers, this pattern is consistent with LS emerging through processing subsequent to the initial feedforward input.

The strong asymmetry of LS between shadow regions and non-shadow regions indicates that the neural computation of lightness is not simply a transformation of local luminance ratios, but instead it depends on the larger-scale interpretation of illumination in which the surface is embedded. This result is consistent with anchoring theory, a prominent mid-level lightness theory (*20*). Anchoring theory proposes that lightness is computed within regions of shared illumination, i.e. by normalizing (anchoring) each luminance value to the highest luminance in the same illumination region, which is perceived as white. For example, **Fig. 1C** contains two illumination regions (a shadow region and a lit region). Square 2 in **Fig. 1C** has the highest luminance within the shadow region, and would thus be perceived as white (normalized to its own luminance). In contrast, square 1 in **Fig. 1C** is the lowest luminance in its illumination region, and would thus be perceived as substantially darker (normalized to the luminance of the square adjacent to square 1, the highest luminance in the lit region). **Fig. 1D** contains a single illumination region, and therefore both square 1 and square 2 would be normalized to the same value, the luminance of the square adjacent to square 1. Hence squares 1 and 2 appear similar in lightness. A prediction of this theory is thus that the lightness of square 1 should differ much less between **Fig. 1C** and **Fig. 1D** than the lightness of square 2, because the normalization computation for square 1 is the same in both scenes, as opposed to that of square 2. This is exactly what we found: the lightness signal dominates in the shadow region (**Fig. 2I**).

Our data help to reconcile low-level and high-level accounts of lightness. Local spatial interactions are clearly capable of generating contextual luminance signals, and such signals are represented early in the visual pathway (*26–30*, *53*). However, our results suggest that these neural populations do not represent surface lightness. Instead, they suggest that the results of early contextual processing are input to a downstream computation that incorporates scene-level illumination structure. In this sense, low-level and high-level mechanisms may represent successive stages of the same inference rather than competing explanations. Thus, even the seemingly elementary distinction between black, gray, and white surfaces is not simply a property of low-level computations early in the visual system, but the outcome of hierarchical cortical inference.

## Acknowledgments

We thank D. Reedy for excellent animal care.

## Author contributions

Conceptualization: FD, PC, TPF

Methodology: FD, PC, TPF

Investigation: FD, RV, TPF

Visualization: FD, TPF

Funding acquisition: RV, TPF

Project administration: TPF

Supervision: TPF

Writing – original draft: FD, TPF

Writing – review & editing: FD, PC, TPF

## Data, code, and materials availability

All data and code used in the analysis will be made available on Figshare and Github upon publication.

## Supplementary Materials

### Materials and Methods

#### Animals

Two adult male rhesus macaques (Macaca mulatta), both aged 12y, were used in this study. The animals were pair-housed when possible in the animal facility at Washington University in St. Louis, and had access to food 24 hours per day in their home cages. On testing days, the animals were placed on water control and earned fluid rewards by performing the tasks. On non-testing days, the animals had ad libitum access to water. All experimental procedures conform to the Guide for the Care and Use of Laboratory Animals and were approved by the Institutional Animal Care and Use Committee (IACUC) of Washington University in St. Louis (protocol 22-0184).

#### Surgery

A titanium recording chamber (inner diameter 21 mm) was implanted over a craniotomy over dorsal Area V4 according to previously published procedures (*37*, *38*). We used a Brainsight Vet robot for neuronavigation (Rogue Research, Montreal, QC, Canada), guided by each animal’s CT and MRI scans. In a separate procedure, the native dura in the chamber was removed and replaced with a silicone artificial dura (*37*, *38*, *54*).

#### Electrophysiology

At the beginning of a recording session, a sterile insert with a thin perforated membrane was positioned in the recording chamber on top of the artificial dura. This insert was used to reduce brain pulsations at the recording site and to facilitate penetration of the artificial dura by applying localized pressure. A Neuropixels 1.0-NHP 10-mm probe (IMEC Inc., Belgium) was then lowered through the artificial dura into the cortex of dorsal Area V4 with a hydraulic microdrive (model MO-972A-oil; Narishige International USA, Inc., Amityville, NY). Voltage signals were continuously monitored, and the probe was advanced until multiunit activity was visible on the deepest ∼3000 µm of the probe. Then, the probe was retracted typically by several hundred µm to ease dimpling of the cortex. Recording sessions began approximately 45 minutes after retraction to allow cortical tissue to recover from dimpling and ensure probe stability. Probe position was varied between recording sessions [RF eccentricity typically between 2 and 10 dva (degrees of visual angle)]. Voltage signals were recorded from the electrode sites positioned in the cortex (distal bank, 384 channels) using a Neuropixels PXIe acquisition module (PXIe_1000; IMEC Inc., Belgium) and SpikeGLX software (https://billkarsh.github.io/SpikeGLX/) using a Windows computer.

Different files recorded during the same recording session were concatenated in temporal order to create a single continuous dataset. Spike sorting was then performed on the concatenated file to extract single- and multi-unit activity using Kilosort4 (*55*). We included all units labeled as ‘good’ by Kilosort in the analyses.

#### Stimulus presentation and task control

During experiments, macaques were seated in a primate chair while they viewed stimuli that were presented onto a rear-projection screen (Aeroview 70; VPixx Technologies Inc., Saint-Bruno, QC, Canada) using an LED projector (PROPixx; VPixx Technologies Inc., Saint-Bruno, QC, Canada). In a minority of sessions, we used a 27-inch Dell LCD monitor (resolution 3840 x 2160 pixels, 60 Hz refresh rate). Display luminance was measured using an X-Rite I1 Display Colorimeter (VPixx Technologies Inc., Saint-Bruno, QC, Canada). The display was positioned 51 cm from the animal’s eyes. Eye position was monitored using an infrared eye-tracking system (EyeLink 1000 Plus; SR Research Ltd., Ottawa, ON, Canada). A photodiode was used to measure when the stimulus appeared on the screen. Stimulus presentation and task were controlled using MonkeyLogic software (https://monkeylogic.nimh.nih.gov/) (*56*). Task and behavioral data (eye position, timestamps of task-related events) were recorded using a data acquisition system (RHD recording controller and I/O Expander; Intan Technologies, Los Angeles, CA). To be able to synchronize neural and behavioral data streams, 32-bit TTL barcodes, generated using an Arduino Uno R3 and sync-barcodes (https://github.com/open-ephys/sync-barcodes), were simultaneously sent to the Neuropixels acquisition system and the Intan RHD acquisition system every 30 seconds.

#### Receptive field mapping

Classical receptive fields (cRFs) were mapped using the subspace reverse correlation approach that we have used in prior studies (*37*, *38*, *49*). Static Gaussian-windowed square-wave gratings (luminance contrast 80%; Gaussian window with full width at half maximum of 2 dva; spatial frequency and phase such that one contrast edge was visible, positioned centrally in the window) and dark gray rings (luminance contrast 80%, diameter 2 dva, ring thickness 0.25 dva) were flashed at random locations selected from a 25 × 25 dva grid with 1 dva spacing, centered at the appropriate visual quadrant. Each stimulus was presented for 50-60 ms while the monkey maintained fixation on a small target. We presented 30 stimuli per trial. Animals received a fluid reward for maintaining fixation throughout the trial. Trials were aborted if eye position deviated from the fixation point. During the recording session, the aggregate cRF of the recorded units was estimated by analyzing high-gamma filtered local field potentials, to properly size and position the lightness stimuli and the stimuli for current-source density analysis.

#### Layer assignment

To determine the laminar position of electrode contacts along the probe, we used the same current-source density (CSD) mapping procedure as in prior studies (*37*, *38*). Briefly, small dark gray rings (94% luminance contrast, 42 ms stimulus duration, ∼700 ms interstimulus interval) were flashed within the cRFs (∼100 stimuli). Current source density profiles were computed as the second spatial derivative of the local field potential across depth. The position of the granular (input) layer was identified as that of the current sink with the shortest latency, following prior studies (*37–39*, *46–51*).

#### Lightness stimuli

We used three main types of scenes: shadow scenes, paint scenes and shuffled scenes. To generate these scenes, we first generated a checkerboard image with alternating dark and light squares (4 rows x 5 columns). The checkerboard was sized such that the cRFs could fit within one square (typical square size was 8×8 dva). This base image thus consisted of squares with two pixel values. Base images differed in the pixel values of both sets of squares keeping the average constant. To generate the shadow scenes, we then multiplied each base image with an illumination matrix. The illumination matrix had the same dimensions as the checkerboard image, and contained two regions, a shadow region and an illuminated region. Both regions were separated by a blurry diagonal border. The values in the illumination matrix were defined by a normal cumulative distribution function, with as argument twice the distance from the diagonal, σ = 1, and scaled such that values in the shadow region were five times lower than those in the illuminated region. By convention we refer to the squares at row 2, column 4, and at row 3, column 2 as squares 1 and 2, where square 2 is the square in the shadow region (**Fig. 1C**). A paint scene was derived from each shadow scene by replacing all pixels in each square with the average value of that square in the shadow scene. A shuffled scene was derived from each paint scene, by randomly permuting the position of all squares except for squares 1 and 2. The luminance of the square with the cRF was thus identical for shadow, paint and shuffled versions of a particular scene. The base image pixel values correspond to those obtained after canceling out the illumination pattern, hence we refer to these as reflectance.

#### Fixation task

Macaques were trained to maintain fixation within 1.2 degrees of visual angle (dva) from a small (0.25 x 0.25 dva) fixation target. In each trial, five randomly chosen lightness scenes were shown. The scenes were positioned and scaled such that the cRFs fell within one square (usually square 2; for the comparisons between dark and light regions (**Fig. 2I,J**), cRFs fell either in square 2 or square 1 by rotating the scenes 180°). Each scene was presented for 500-800 ms. In most sessions successive scenes were interleaved with a blank screen, which was presented for 200 ms. Each blank screen had the same average luminance as that of the scene that followed it.

#### Macaque psychophysics: stimuli, task and analysis

We used similar shadow and paint scenes as described above, but with a pseudorandom pattern of square luminance values (**Fig. 1E,F**). This was done to avoid that the animal would compare two neighboring squares rather than squares 1 and 2. This pattern differed for scenes with different luminance contrast between the squares to be compared (see below). To generate these scenes, we started with a base image with a shuffled distribution of the reflectance values for each square. Squares were 8 dva wide. In three sessions of monkey B, we used scenes in which the central column of the scene consisted of rectangles (2 dva width x 8 dva height). The difference between shadow and paint data was similar when excluding those sessions. Each half of the scene consisted of the same set of reflectance values (excluding the squares that the monkey had to compare, see below). The reflectance values for the four squares adjacent to the illumination border were chosen from the same set for each half of the board, in random order (see below for the corresponding luminance values). The reflectance values for the five remaining squares that bordered the edge of the scene were also chosen from the same set for each half, in random order. The two remaining squares are squares 1 and 2, which the monkey was trained to compare. Reflectance values for squares 1 and 2 were set to different values in different versions of the base image, such that their average remained the same (typically 18 pairs of reflectance values, equally divided between conditions in which square 2 had higher reflectance than square 1 and vice versa). As described above, we then applied the illumination pattern to these base images to generate shadow scenes (illumination matrix defined by a normal cumulative distribution function as described above). Either the lower left or the upper right half of the scene was covered by shadow. For each of these shadow scenes, we derived a corresponding paint scene, using the procedure described above. In two sessions from monkey B, squares 1 and 2 were labeled. Labels were (0.38 dva x 0.38 dva) square reflectance markers in each of the squares that neighbored squares 1 and 2 (reflectance values for these markers were chosen from the same set for both squares). The difference between shadow and paint data was similar between sessions with or without labels. For each condition, we generated several versions (typically five), which differed in the pattern of reflectance values for the other squares. Square 2 had typically one of these luminance values: [2.5; 8.8; 15.0; 21.4; 27.7; 40.4; 60.0; 85.3; 111.0; 123.9; 143.3; 169.1; 188.5; 201.5; 208.0; 214.5; 221.0; 227.4] cd/m^2^.

Each trial began with the presentation of a fixation point on a gray background. Monkeys were required to acquire and maintain fixation on a small (typically 0.26 dva x 0.26 dva) fixation target during 300 ms (typically within 1.5 dva). Then a shadow or paint scene was presented for 800-2000 ms, on a black background. The scene was positioned such that the fixation point was placed at the center of the scene, thus equidistant from the center of each of squares 1 and 2. At the end of the trial the fixation point disappeared, cueing the animal to indicate which of these squares appeared lighter by making a saccade to one of two choice targets that appeared at the locations of these squares.

Conditions with luminance values to the left from the vertical dashed line, or to the right from the vertical dotted line in **Fig. 1G** are unambiguous: the animal’s choice would be the same irrespective of whether it reported the sign of luminance or reflectance difference between both squares. Conditions with luminance in between the vertical dashed/dotted lines in **Fig. 1G** are ambiguous: the choice would be different depending on whether the animal relied on lightness or luminance. Similar to other NHP studies that use illusions (*57*) we rewarded animal B veridically (i.e. according to the luminance difference between the squares) only for the unambiguous conditions, and randomly for the ambiguous conditions. To further minimize the possibility that the animal would adjust its strategy for ambiguous conditions, these were only shown in up to 20% of trials. We wondered if the effect required this specific reward paradigm. We therefore rewarded animal T veridically, i.e. according to the sign of the luminance difference between the squares, for all conditions. We found a significant difference between shadow and paint scenes in both animals (**Fig. 1I**).

For each scene type (shadow and paint scenes), we computed the proportion of trials in which the monkey chose square 2 rather than square 1 as the one that appeared higher in luminance. We quantified lightness constancy using the Thouless ratio (*τ*) (*8*, *35*, *36*). *τ* is defined as

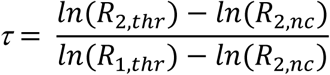

where *R*_2,*t*ℎ*r*_ and *R*_1,*t*ℎ*r*_ are the reflectance values of squares 2 and 1 at the point where the psychometric function reaches 0.5 (i.e., the point of subjective equality), and *R*_2,*nc*_ is the reflectance that square 2 would have at the point of subjective equality in the absence of lightness constancy (i.e., the condition in which squares 1 and 2 have identical luminance).

Thouless ratio τ = 1 indicates perfect lightness constancy (perfect report of reflectance), whereas τ = 0 indicates absent lightness constancy (perfect report of luminance). We fitted a logistic function *f(x)* through the data for each scene type and each session using the MATLAB function ‘fit’ with fit method non-linear least squares.

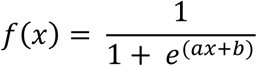

Lower bounds, upper bounds and start point for parameters *a* and *b* were respectively [-Inf -Inf], [0 Inf] and [-5 0]. We required that the fraction of choices for the lowest and highest luminance values of square 2 were respectively <0.1 and >0.9, and that R^2^ of the logistic function fits for both shadow and paint conditions was >0.9 for the analysis in **Fig. 1I**.

To statistically compare the psychometric functions of shadow and paint scenes, we used a logistic binomial model (MATLAB function ‘fitglm’ with parameter ‘Distribution’ set to ‘binomial’). To determine whether *τ* for paint was significantly different from 0, we used bootstrapping to compute the confidence interval (MATLAB function ‘bootci’, 2000 bootstraps). To compare *τ* for shadow and *τ* for paint across sessions, we used a paired t-test.

#### Human psychophysics: stimuli, task and analysis

The psychophysics experiments in human subjects were approved by the Institutional Review Board (IRB) at Washington University in Saint Louis (project ID 202311107). Participants were compensated for their time. We conducted the experiment in naïve healthy adult subjects (n=10, 5 female and 5 male, ages between 23 and 27 years old). During the task, participants were seated with their head in a chinrest, with their eyes at a distance of 63.5 cm from the monitor on which the stimuli were shown (27-inch Dell LCD monitor, resolution 3840 x 2160 pixels, 60 Hz refresh rate). Their eye position was tracked with an infrared eye-tracking system (EyeLink 1000; SR Research Ltd., Ottawa, ON, Canada). Subjects conducted a similar psychophysical task that we used in macaques. Subjects were instructed to report which one of two labeled squares in a lightness scene was closest in appearance to a white square. Shadow and paint scenes were constructed from a 4×5 base image with pseudorandom pattern of reflectance values, as explained above. Squares were 5.5 dva wide. The two squares to be compared (squares of interest: row 3, column 2, and row 2, column 4) were labeled by four small (width 0.13 dva) square markers positioned in the neighboring squares. Each marker had the same luminance as the square on the opposite side from the square of interest, relative to the square in which the marker was positioned. The square of interest in the dark region (square 2) had one of these luminance values: [0.5; 4.1; 6.0; 8.1; 10.6; 12.0; 12.3; 14.2; 16.6; 18.8; 20.6; 24.1] cd/m^2^. The set of luminance values of the other squares in the shadow region was typically [1.1; 3.3; 3.9; 9.6; 9.9; 18.3; 21.1; 31.3; 37.6] cd/m^2^. The luminance of the squares in the non-shadow region was five times higher than those in the shadow region.

Subjects were required to fixate a small (0.26 dva x 0.26 dva) target throughout each trial. Five hundred ms after acquiring fixation, a lightness scene was presented for 500 ms, positioned such that the fixation appeared in the center of the scene, on a black background. Shadow and paint scenes were randomly interleaved. After the scene disappeared, a blank screen appeared for 700 ms. Then the fixation point disappeared and subjects were required to indicate their response with a button press. Subjects received visual feedback about which square they chose (but not about the correct answer). Trials were aborted if subjects broke fixation (radius 1.5 dva). These data were analyzed similarly to the behavioral data obtained in macaques.

#### Between-condition decoding analyses

We performed between-condition decoding analyses using random forest regressors to compare the difference in luminance representation between paint scenes and shadow scenes (for the lightness signal, LS) and between paint scenes and shuffled scenes (for the simultaneous contrast signal, SCS). To train the random forest models, we used the TreeBagger function in MATLAB with options Method=’regression’, and 200 trees. Models were trained to predict the luminance of the square on which the cRFs were positioned from spike counts evoked during the stimulus presentation (summed between 0 and 500 ms after stimulus onset). We used data from 90% of the trials from one scene type (e.g. paint, in case of LS) to train the model. Each unit was a predictor, and each trial an observation. After training the model, we then tested it on all trials from the other scene type (e.g. shadow), and also on the 10% left-out trials from the scene type used to train the model. LS was defined as the average difference between predicted luminance from shadow scenes and paint scenes (across decoder cycles and luminance values), and SCS was defined as the average difference between predicted luminance from paint scenes and shuffled scenes (across decoder cycles and luminance values). To statistically compare predicted luminance in the between-condition decoding analyses between different test conditions we used a permutation test (shuffling scene labels): a null distribution for LS (or SCS) was created by shuffling the scene type labels 2000 times, and the p value was estimated as the fraction of this distribution for which LS (or SCS) was at least as large as the measured value. We used one-sample t-tests to determine whether the distribution of LS and SCS across penetrations was significantly larger than 0.

#### Laminar analysis of LS and SCS

To compare LS and SCS between laminar compartments, we generated pseudopopulations of units from the sessions for which laminar compartments could be identified from the CSD analysis (see above). Because the number of units differed between laminar compartments we subsampled the data. In each decoder cycle, we subsampled a fixed number of units for each laminar compartment (n = number of units in granular layer), with replacement, and a fixed number of trials for each unit (t = number of trials available for that combination of luminance and scene type), with replacement. LS or SCS was then computed within each compartment using the approach above. LS (SCS) was averaged across 150 decoding cycles.

#### Dynamics of contextual information

To characterize the temporal dynamics of contextual information, we used demixed principal component analysis (dPCA)(*52*). Only sessions in which a blank scene was presented between successive scenes were included, to minimize interference from the preceding stimulus. Neural firing rates were calculated in non-overlapping 1-ms bins and smoothed by convolution with a Gaussian kernel (20-ms width). Neural activity was analyzed over a time window extending from 100 ms before to 500 ms after stimulus onset. We generated bootstrapped pseudo-populations (1,000 bootstrap iterations). In each iteration, neurons were randomly sampled with replacement. To balance trial numbers across recording sessions, we determined the minimum number of available trials for each luminance level across sessions. Within each bootstrap iteration, this number of trials was randomly selected without replacement from each session, and the selected trials were pooled across sessions. Condition-averaged population responses were then used to calculate the marginalizations and perform dPCA. The regularization parameter (λ) was optimized once using the dpca_optimizeLambda function provided in the dPCA MATLAB toolbox (*52*), using the full neuronal population, and was held constant across all bootstrap iterations.

Luminance and scene type (shadow or paint, or paint or shuffled) were used as independent variables. We focused on the first component associated with the scene type marginalization, which captured the largest amount of scene type related variance.

To estimate the latency of contextual signals, we calculated the absolute difference between the time-varying trajectories of the first scene-type dPCA component for the two scene types in each analysis. Latency was defined as the earliest time point at which this difference exceeded 33% of its maximum value and remained above this threshold for at least 5 consecutive milliseconds. Latency was calculated independently for each bootstrap iteration, yielding 1,000 latency estimates.

#### Dynamics of LS and SCS

After training random forest regressors to predict luminance from the spike rate in a window [0 500] ms after stimulus onset, we tested these decoders in 100-ms sliding windows (1 ms step) to characterize the dynamics of LS and SCS (**Fig. 4C**). LS and SCS were computed as above in each window. The statistical significance of each signal in each window was assessed by subtracting the baseline value (from windows prior to stimulus onset, starting at −125 ms) and the p-value computed by comparing the result with the highest amplitude obtained after shuffling the scene type labels. Latency was defined as the first time point for which p<0.05 for 15 adjacent windows.

#### Neural lightness constancy

To quantify neural lightness constancy, we computed neural Thouless ratios. These were derived after excluding neurons in the input layer, as our laminar analysis only found LS in extragranular compartments. We thus used the sessions with identifiable laminar compartments, for which there was a significant LS and SCS signal. For each session, we trained a random forest regressor using neuronal responses to shuffled scenes across the full range of luminance values. The decoder was trained to predict the luminance of the square with the cRFs (square 2) relative to that of square 1. The trained decoder was then tested on neuronal responses to shadow scenes for which the cRFs were positioned in the dark region of each scene. For each luminance contrast, we calculated the percentage of decoder predictions for which the luminance of square 2 is predicted to be higher than that of square 1. From these data we then derived neural Thouless ratios, using the same procedure as for the behavioral data.

---

**Fig. S1.**
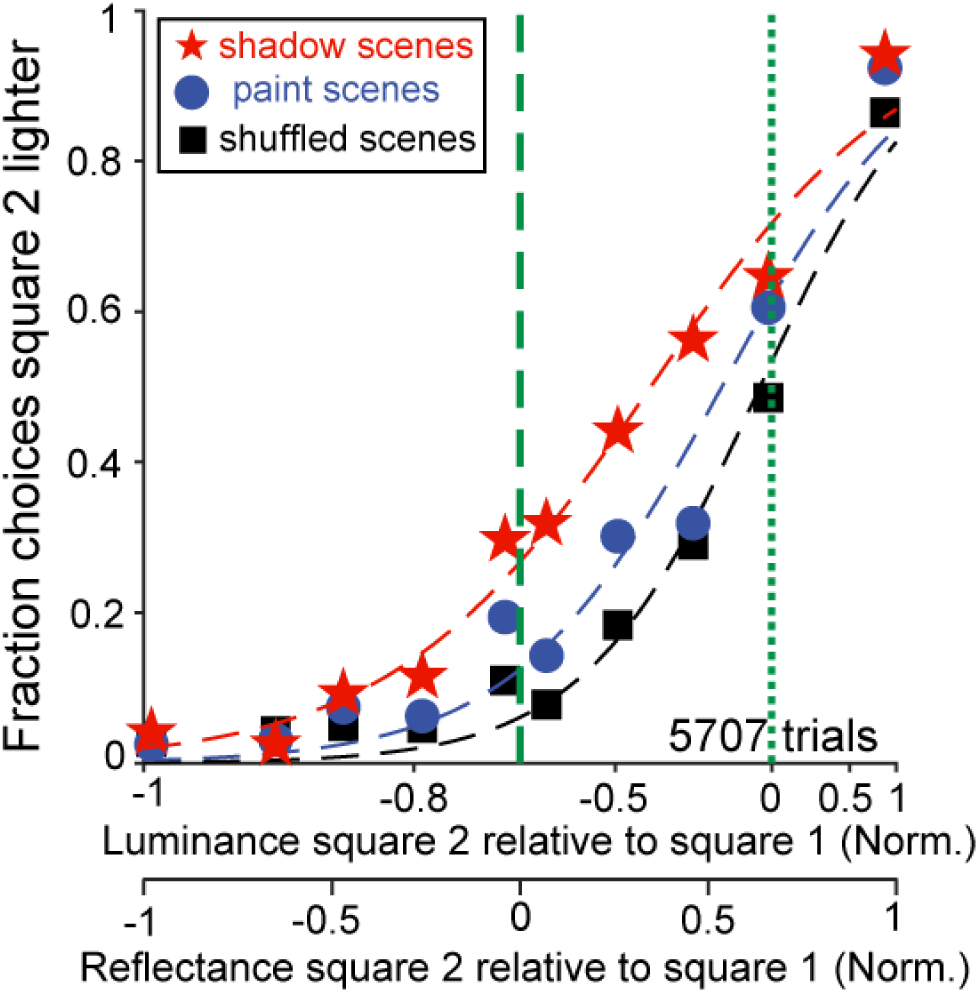
Behavioral data for shadow, paint and shuffled scenes. Similar to Fig. 1G, for three behavioral sessions in monkey B in which we included shuffled scenes. These scenes are derived from paint scenes by randomly repositioning all squares except for the two squares that the animal is comparing. The data from shadow scenes is significantly shifted relative to those of paint scenes (logistic regression model shadow and paint data, p-value for scene type: p = 4.6×10^-5^), and the data from paint scenes is significantly shifted relative to those of shuffled scenes (logistic regression model paint and shuffled data, p-value for scene type: p = 1.2×10^-9^).

**Fig. S2.**
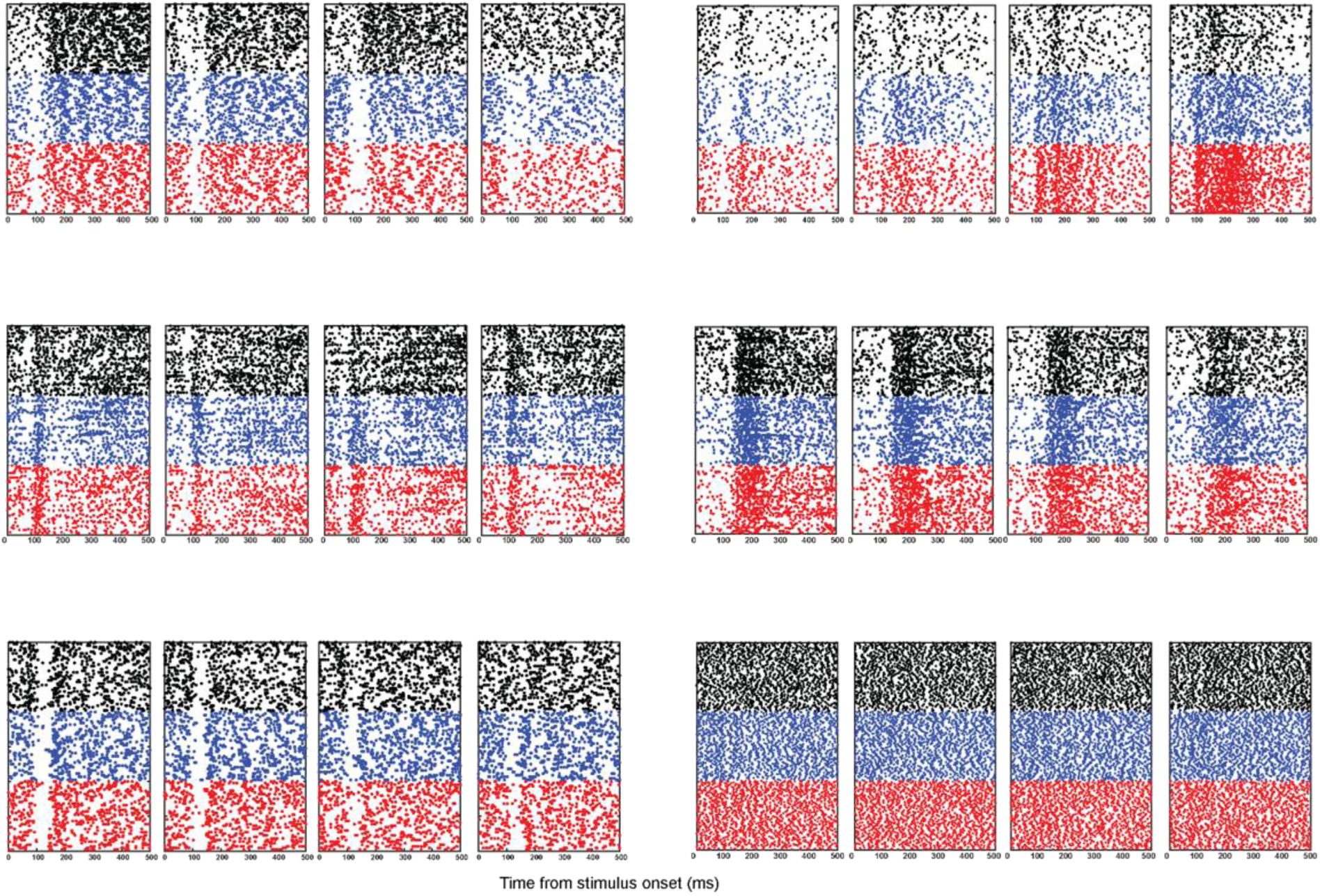
Dot rasters for example units. Each dot shows a spike. Red, blue and black indicate respectively data from shadow, paint and shuffled scenes. Each of the panels shows data for a different luminance. Units correspond to those in Fig. 2D.

**Fig. S3.**
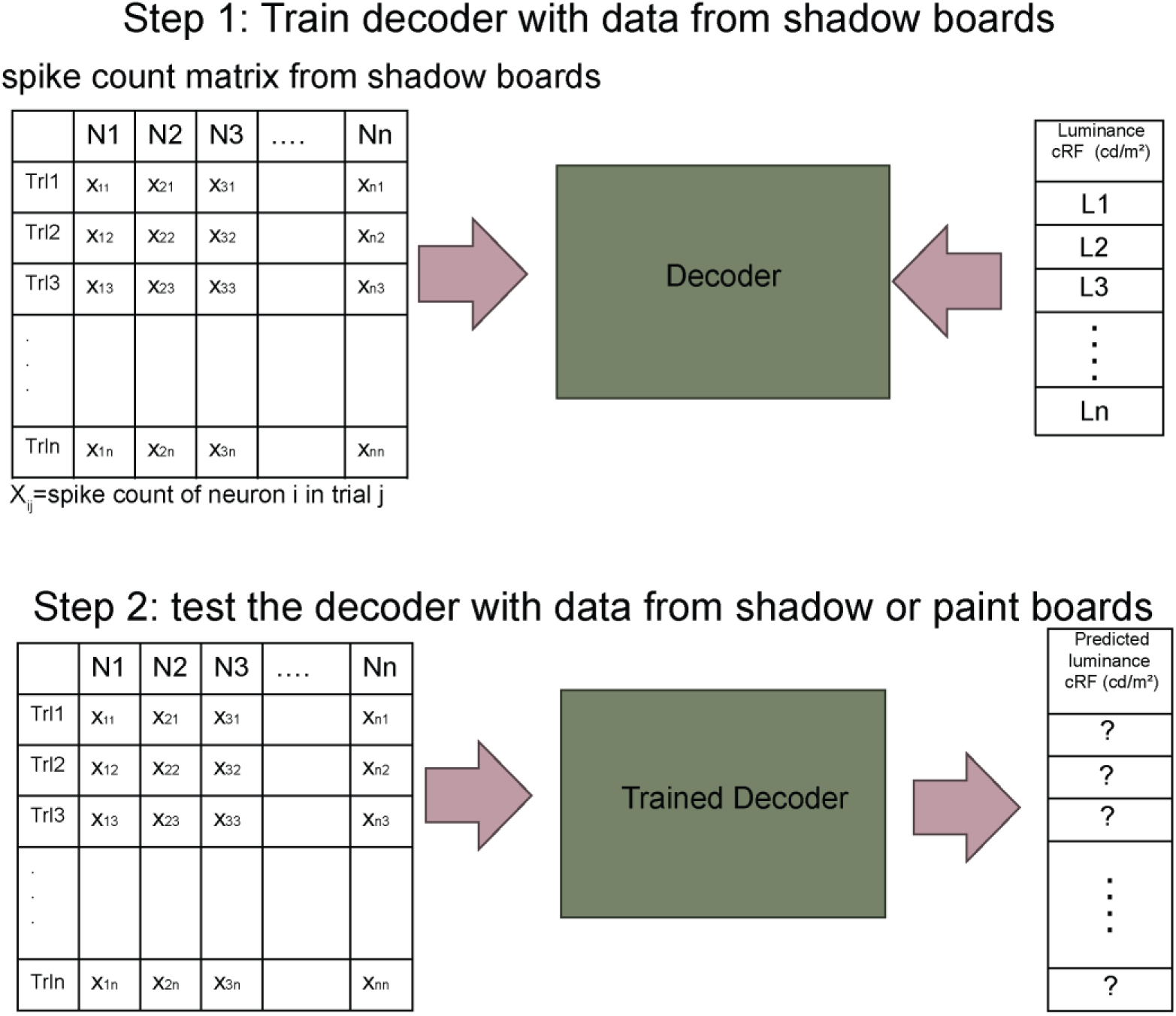
Schematic of between-condition decoding analysis.

**Fig. S4.**
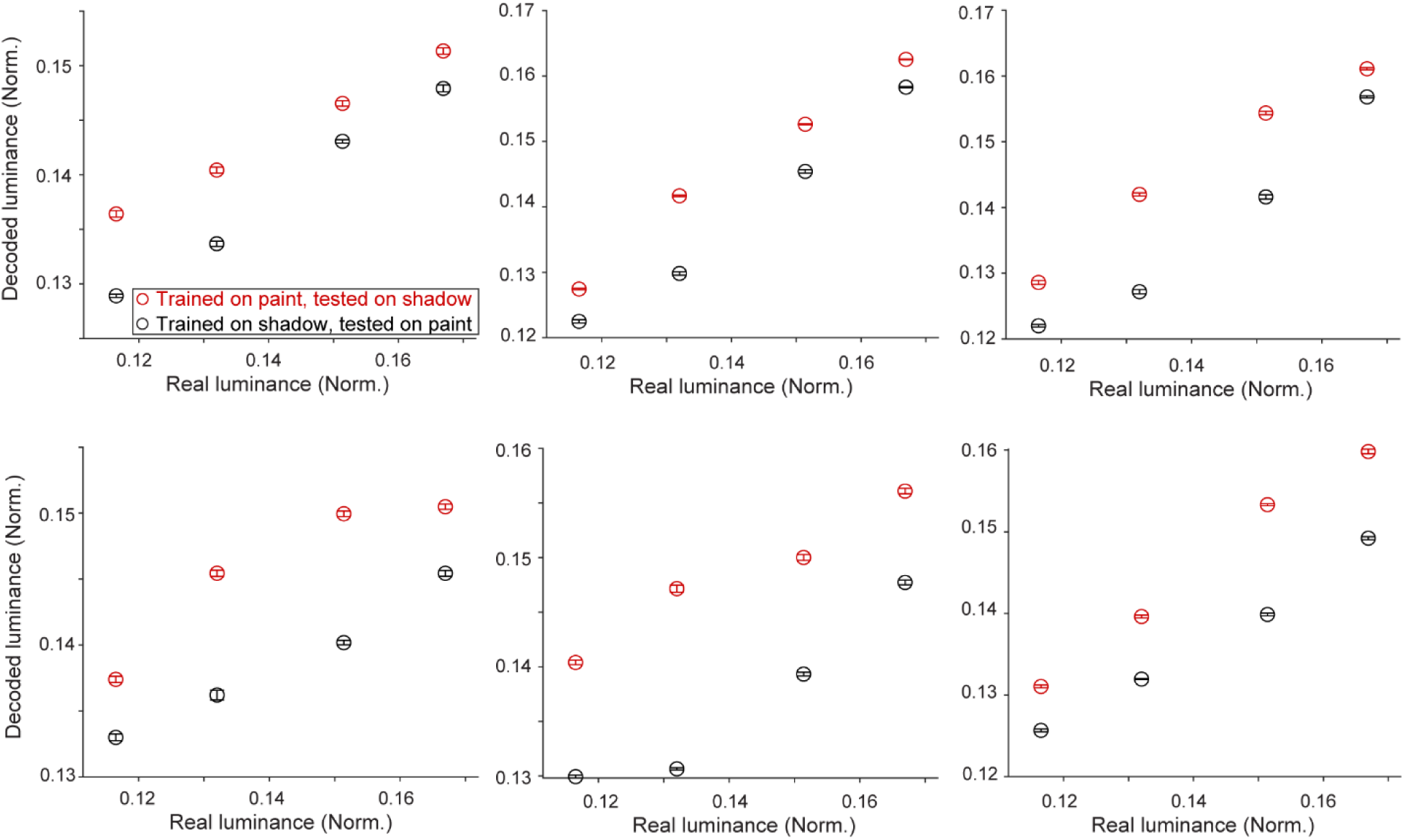
Lightness bias is also apparent when training decoders on data from one scene type and testing them on data from the other scene type. Random forest regressors were trained on paint (or shadow) scene responses to decode the luminance in the cRFs. After training, decoders were tested on data from the alternative configuration, i.e. shadow (or paint) scene responses, recorded during the same penetration. Symbols indicate mean ± standard error of the mean across five decoder runs. Each panel shows data from a different penetration. Top row: data from monkey B. Bottom row: data from monkey T. In each panel, the LS is statistically significant (permutation test p < 0.05).

**Fig. S5.**
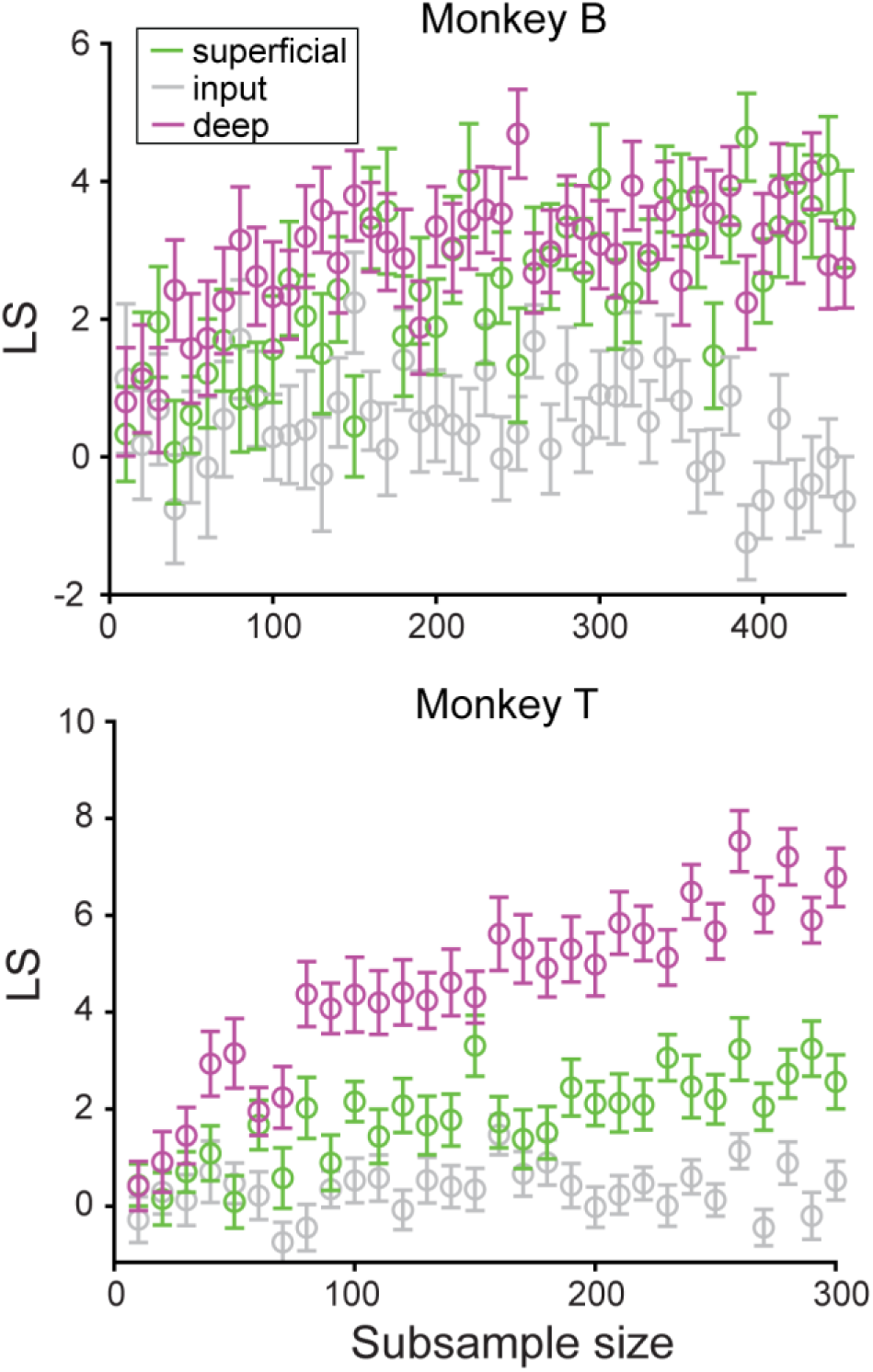
Lightness signal is weak in input layer irrespective of subsample size. Mean ± standard error of the mean across 50 decoder cycles per subsample size.

